# Zinc detoxification: a functional genomics and transcriptomics analysis in *Drosophila melanogaster* cultured cells

**DOI:** 10.1101/179598

**Authors:** Stephanie E. Mohr, Kirstin Rudd, Yanhui Hu, Wei Roc Song, Quentin Gilly, Michael Buckner, Benjamin E. Housden, Colleen Kelley, Jonathan Zirin, Rong Tao, Gabriel Amador, Katarzyna Sierzputowska, Aram Comjean, Norbert Perrimon

## Abstract

Cells require some metals, such as zinc and manganese, but excess levels of these metals can be toxic. As a result, cells have evolved complex mechanisms for maintaining metal homeostasis and surviving metal intoxication. Here, we present the results of a large-scale functional genomic screen in *Drosophila* cultured cells for modifiers of zinc chloride toxicity, together with transcriptomics data for wildtype or genetically zinc-sensitized cells challenged with mild zinc chloride supplementation. Altogether, we identified 47 genes for which knockdown conferred sensitivity or resistance to toxic zinc or manganese chloride treatment, and more than 1800 putative zinc-responsive genes. Analysis of the ‘omics data points to the relevance of ion transporters, glutathione-related factors, and conserved disease-associated genes in zinc detoxification. Specific genes identified in the zinc screen include orthologs of human disease-associated genes CTNS, PTPRN (also known as IA-2), and ATP13A2 (also known as PARK9). We show that knockdown of *red dog mine (rdog; CG11897)*, a candidate zinc detoxification gene encoding an ABCC-type transporter family protein related to yeast cadmium factor (YCF1), confers sensitivity to zinc intoxication in cultured cells and that *rdog* is transcriptionally up-regulated in response to zinc stress. As there are many links between the biology of zinc and other metals and human health, the ‘omics datasets presented here provide a resource that will allow researchers to explore metal biology in the context of diverse health-relevant processes.

## INTRODUCTION

Whereas metals such as mercury or cadmium are solely toxic to cells, other metals, such as zinc and manganese, are essential for cell viability and toxic only in excess. Zinc is a structural component of many proteins and is also thought to act as a signaling molecule (Fukada *et al.* 2011). Adding to the complexity of cellular zinc regulation, zinc is maintained at different levels in different organelles. Cells have evolved complex mechanisms for surviving zinc insufficiency, maintaining cellular and subcellular zinc homeostasis, and surviving exposure to toxic levels of zinc. The molecular mechanisms underlying regulation of zinc homeostasis and detoxification are in some cases zinc-specific and in other cases, relevant to other metals. Methods used by cells to maintain zinc levels and/or survive metal toxicity include regulation of proteins required for metal influx (e.g. ZIP family importers of zinc), metal efflux (e.g. ZnT family exporters of zinc), or metal chelation (e.g. by metallothionines that chelate metals), as well as sequestration of zinc and/or biomolecules damaged by zinc in membrane-bound organelles such as the yeast vacuole or mammalian lysosome (Kambe *et al.* 2015).

In addition to involving transporters and chelators, metal detoxification also involves more general detoxification strategies. Glutathione (GSH) has long been known to have the ability to form a complex with zinc or cadmium (Perrin and Watt 1971). Data from plants, yeasts, tunicates, fish, and other organisms suggest the relevance of GSH levels, conjugation, and/or transport to metal detoxification (Perego and Howell 1997; Penninckx 2002; Gharieb and Gadd 2004; Franchi *et al.* 2012; Seth *et al.* 2012). Genetic evidence provides further support for a connection between GSH and metal detoxification. The ABCC-family transporter yeast cadmium factor (YCF1) is one example. YCF1 was originally identified based on a cadmium sensitivity phenotype and has been implicated in GSH-mediated detoxification of cadmium. The YCF1 protein is thought to be localized to the vacuole and to mediate transport of bis(glutathionato)cadmium into the vacuole (Nagy *et al.* 2006). Evidence suggests that the related protein YOR1, which is localized to the plasma membrane, can transport GSH-conjugated cadmium out of the cell. Consistent with this idea, *ycf1, yor1* double mutant yeast strains are reportedly more sensitive to cadmium intoxication than either single mutant strain (Nagy *et al.* 2006).

The biology of zinc and other metals has many connections with human diseases. For example, genetic disruption of genes encoding metal transporters can lead to diseases of metal insufficiency or excess (Clayton 2017). In addition, accumulation of high levels of metals, which can occur following consumption of contaminated drinking water or through occupational exposure, can negatively impact human development and cause disease. Further, because metals are used by cells as ‘weapons’ in defense against pathogens (Weiss and Carver 2017), metal insufficiency can impact immune function. Moreover, metals or metal-related genes have been implicated in, or levels correlated with, diseases such as diabetes, Parkinson’s disease, and Alzheimer’s disease (Chasapis *et al.* 2012). The zinc transporter ZNT8 is a common autoantigen in type 1 diabetes (Arvan *et al.* 2012), for example, and ATP13A2 (PARK9), a Parkinson’s disease gene, has been implicated in zinc homeostasis (Kong *et al.* 2014; Park *et al.* 2014; Tsunemi and Krainc 2014). Furthermore, adaptation of insect vectors of disease such as mosquitoes to metals might confer concomitant resistance to insecticides (Poupardin *et al.* 2008), such that understanding metal detoxification in insects might impact our understanding of disease vector control and the impact of polluted environments on the spread of insect-borne diseases (Poupardin *et al.* 2012).

Although yeast provides an excellent genetic platform for study of the cell biology of metal detoxification, using a single-celled organism has limited potential to model multicellular systems such as humans or insect vectors of disease. *Drosophila* presents many advantages as a genetic model system, including to study metal homeostasis and detoxification at the cellular and whole-organism levels; to study the effects of genetic or environmental perturbation of metal levels in the context models of human diseases (e.g. in *Drosophila* models of Parkinson’s or Alzheimer’s disease); and as a model of adaptation to metals and/or insecticides by insect vectors of disease. Work by several labs has established *Drosophila* as an *in vivo* model for study of zinc biology in a multicellular system (Richards and Burke 2016; Xiao and Zhou 2016), as well as for evolutionary studies of metal-related genes (Sadraie and Missirlis 2011; Rempoulakis *et al.* 2014). R. Burke and colleagues in particular performed a comprehensive genetic survey of *in vivo* ZIP and ZnT family zinc transporter functions using combined knockdown and over-expression approaches (Lye *et al.* 2012; Lye *et al.* 2013). Moreover, R. Burke, B. Zhou, and others have established the fly gut as a system for the study of zinc and other metals (Wang *et al.* 2009; Wang and Zhou 2010; Jones *et al.* 2015). Studies in *Drosophila* have also identified a role for zinc in kidney stone disease (Chi *et al.* 2015), and *Drosophila* is an established model in which to study the effects of metal-containing nanomaterials (Alaraby *et al.* 2016).

Altogether, the existing literature suggests that *Drosophila* provides an excellent system in which to study the cellular and organismal biology of metal homeostasis and detoxification. Despite the growing body of work in *Drosophila* on zinc biology and other metal-related studies, however, there has remained a need for the application of high-throughput functional genomic methods to the study of zinc and other metals in *Drosophila*. Here, we describe the results obtained by applying two complementary ‘omics approaches to the identification of genes relevant to metal homeostasis and detoxification. Specifically, we performed large-scale *Drosophila* cell-based RNAi screens to identify genes relevant to zinc or manganese detoxification, and performed a transcriptome-wide analysis of genes regulated in response to mild metal supplementation of wildtype or genetically zinc-sensitized cells. The results point to conserved genes and functions, and provide a resource for further study.

## MATERIALS AND METHODS

### Cultured cell lines

The screen was performed using the DRSC isolate of the S2R+ *Drosophila* cell line. Derivatives of this cell line newly generated in this work are available from the *Drosophila* Genome Resource Center (DGRC) cultured cell repository in Bloomington, IN, USA (DGRC cell IDs 1000 and 1001; see below and Supplemental Reagents Table).

### Cell RNAi screening

We screened in total four dsRNA reagent libraries for *Drosophila* cell-based RNAi screening from our *Drosophila* RNAi Screening Center (DRSC) collection (Hu *et al.* 2017): the TM library targeting genes encoding transmembrane domain-containing protein library (17 unique 384-well assay plates), AUTGY library targeting genes encoding autophagy-related factors (3 plates), MBO1 library targeting genes encoding proteins associated with membrane-bound organelles (2 plates), and a custom-designed plate with candidate metal-related factors we refer to as the “Megadeath” plate (1 plate). In all cases, experimental dsRNAs are excluded from the outermost two wells of the final 384-well assay plate design to limit edge effects. Three replicates of each unique plate in the library (metal-supplemented conditions) or two replicates of each unique plate (control) were screened. To perform the screens, we added S2R+ cultured cells to dsRNAs-containing assay plates as described previously (Echeverri and Perrimon 2006). We then incubated the plates in a 25°C incubator with humidity control for four days. Next, freshly prepared ZnCl_2_ or MnCl_2_ (Sigma Aldrich) in solution or a control treatment (water) was added to the assay plates using a Formulatrix Mantis liquid handling robot to a final level of supplementation of 15 mM. Twenty-four h following metal supplementation or control treatment, cells were lysed and total ATP levels per well were determined using Promega Cell Titer Glo and a Molecular Devices Spectramax Paradigm luminometer. The step-by-step screen and assay protocols we used are available online at <https://fgr.hms.harvard.edu/fly-cell-rnai-384-well-format> and <https://fgr.hms.harvard.edu/fly-cell-total-atp-readout>. Relative luciferase values for each plate were normalized to the plate average, replicates were averaged, and average normalized relative luciferase values were then compared across plates by calculating Z-scores.

### Generation of CRISPR knockout cell lines

The sgRNA sequence used to target *ZnT63C* was TGTGACCAATTCGATGGCTC; the sgRNA sequence used to target IA-2 was CGGCTGTTCCGCGTGCTCTCTGG (see also Supplemental Reagents Table). The sgRNAs were cloned and introduced into cells as described in (Housden *et al.* 2014; Housden *et al.* 2015). Briefly, following introduction of Cas9 and sgRNAs targeting *ZnT63C* or *IA-2* and single-cell isolation, we used high-resolution melt analysis (HRMA) to identify gene-modified cells. The *ZnT63C* or *IA-2* gene regions from colonies positive by HRMA were then amplified by PCR, PCR products were individually cloned by TOPO cloning, at least 10 isolates were subjected to Sanger sequencing, and sequence data was aligned and analyzed to confirm that all alleles contained frameshift mutations.

### RNA preparation and RNAseq analysis

For RNAseq analysis of control or metal-treated cells, wildtype S2R+ or CRISPR modified mutant cell derivatives (*IA2-KO* and *ZnT63C-KO*) were first grown to confluency in 10 mL of media in T-75 flasks; then control samples were left untreated and experimental samples were supplemented to a final concentration of 1 mM ZnCl_2_ or MnCl_2_; and cells were incubated for 24 h, centrifugated, and resuspended in TRIzol reagent (Thermo Fisher). RNA was extracted as described previously using chloroform extraction and isopropanol precipitation (Song *et al.* 2010). Each final total RNA solution was divided into two aliquots, one of which was freshly used for RNA Integrity Number (RIN) evaluation at the Harvard Medical School Biopolymers Facility. If the RIN was >6.7 and no RNA degradation was observed following analysis, the other aliquot (stored at −80°C) was shipped on dry ice to the Columbia Genome Center (Columbia University, New York, NY) for standard sample processing and raw data analysis. The raw data files were processed by the Columbia Genome Center. In order to compare the results at the gene level, we took the average of the FPKM values for each gene for the two replicates done for each condition, then determined the log_2_ ratio of FPKM levels for each genotype and treatment condition combination vs. FPKM levels in the same genotype. Prior to this analysis, we set to a value of “1” any average FPKM value between 0 and 1 to reduce the possibility that we get large ratio values for genes with negligible levels of detected transcript in both the experimental sample and the wildtype control (e.g. the ratio of FPKM 0.1 vs. 0.0001), as we assume those ratios are not likely to have biological relevance. A cutoff of two-fold change for all replicates was applied (log_2_ >1 or <-1).

### qPCR analysis

Wildtype S2R+ cells were cultured under standard growth conditions in a 75 mL flask for 72 h. Next, the appropriate amount of a 1 M stock solution of ZnCl_2_ or ZnSO_4_ (or water) to reach a final supplementation concentration of 0, 1, 3, or 5 mM. Cells were then incubated for 24 h at 25°C, centrifugated and resuspended in TRIzol (Thermo Fisher), treated with DNaseI, and cDNA was prepared using a QIAGEN RNeasy kit. The cDNA was analyzed by real-time quantitative PCR (qPCR) using SYBR Green and with tubulin as an internal reference gene control. All experiments were conducted in triplicate qPCR reactions. Data were analyzed with Bio-Rad CFX Manager software. Additional details regarding reagents and the sequence of oligonucleotide primer sequences used to detect experimental and control genes are indicated in the Supplemental Reagents Table.

### Enrichment analysis of the RNAi screen and RNAseq datasets

The gene hits from RNAi screen and RNA-Seq profiling were analyzed for over-represented gene sets using an in-house JAVA program based on hyper-geometric distribution. The gene sets we queried were assembled using gene ontology annotation, pathway annotation from GLAD (Hu *et al.* 2015), and the *Drosophila* protein complex annotation from COMPLEAT (Vinayagam *et al.* 2013). Human pathway annotation of Reactome (Croft *et al.* 2011) and KEGG (Kanehisa *et al.* 2017) were mapped to *Drosophila* gene sets using DIOPT (Hu *et al.* 2011) and included in enrichment analyses.

### Data availability

High-confidence gene-level hits from the RNAi screens are shown in Fig. 1 and listed with additional details (human orthologs, Z-scores, etc.) in Supplemental Tables 1, 2, and 3. In addition, a complete list of gene-level high-, moderate-, and low-confidence RNAi screen hits, as well as and enrichment analysis results, is provided in Supplementary file 1. Reagent-level RNAi screen data are available from the FlyRNAi database of the DRSC (see “Screen Summary” for a full list of screen datasets or “Gene Lookup” to query by gene or reagent) (Hu *et al.* 2017). The RNAi screens were assigned DRSC Project IDs 177 through 185 and 193. To view the full dataset for a screen, replace “X” with the three-digit DRSC Project ID in the URL <http://www.flyrnai.org/cgi-bin/RNAi_public_screen.pl?project_id=X>. The RNAi screen datasets are also available at NCBI PubChem BioAssay (Wang *et al.* 2014). The screens were assigned PubChem BioAssay IDs 1259314, 1259315, 1259315, 1259316 (additional depositions in process). To view a dataset at PubChem BioAssay, replace “X” with the seven-digit PubChem ID in the URL <https://pubchem.ncbi.nlm.nih.gov/bioassay/X>. Analyzed results of the transcriptomics study are summarized in Table 1, Fig. 2, and Fig. 3, and provided in full in Supplementary file 2 (gene-level data and enrichment analysis results). In addition, the RNAseq datasets are available from NCBI Gene Expression Omnibus (GEO), GEO accession ID GSE99332 (<https://www.ncbi.nlm.nih.gov/geo/query/acc.cgi?acc=GSE99332>).

**Table 1:** Summary of transcriptomics analysis of wildtype and zinc-sensitized cells under control or metal-supplemented conditions.

|  | **wildtype S2R+** | ***IA2-KO*** | ***ZnT63C-KO*** |
| --- | --- | --- | --- |
| + 1 mM ZnCl<sub>2</sub> | 319 down<br>121 up | 66 down<br>128 up | 998 down<br>835 up |
| +1 mM MnCl<sub>2</sub> | 23 down<br>33 up | 26 down<br>26 up | 68 down<br>16 up |

Down, down-regulated as compared with untreated cells of the same genotype; up, up-regulated as compared with untreated cells of the same genotype.

**Figure 1:**
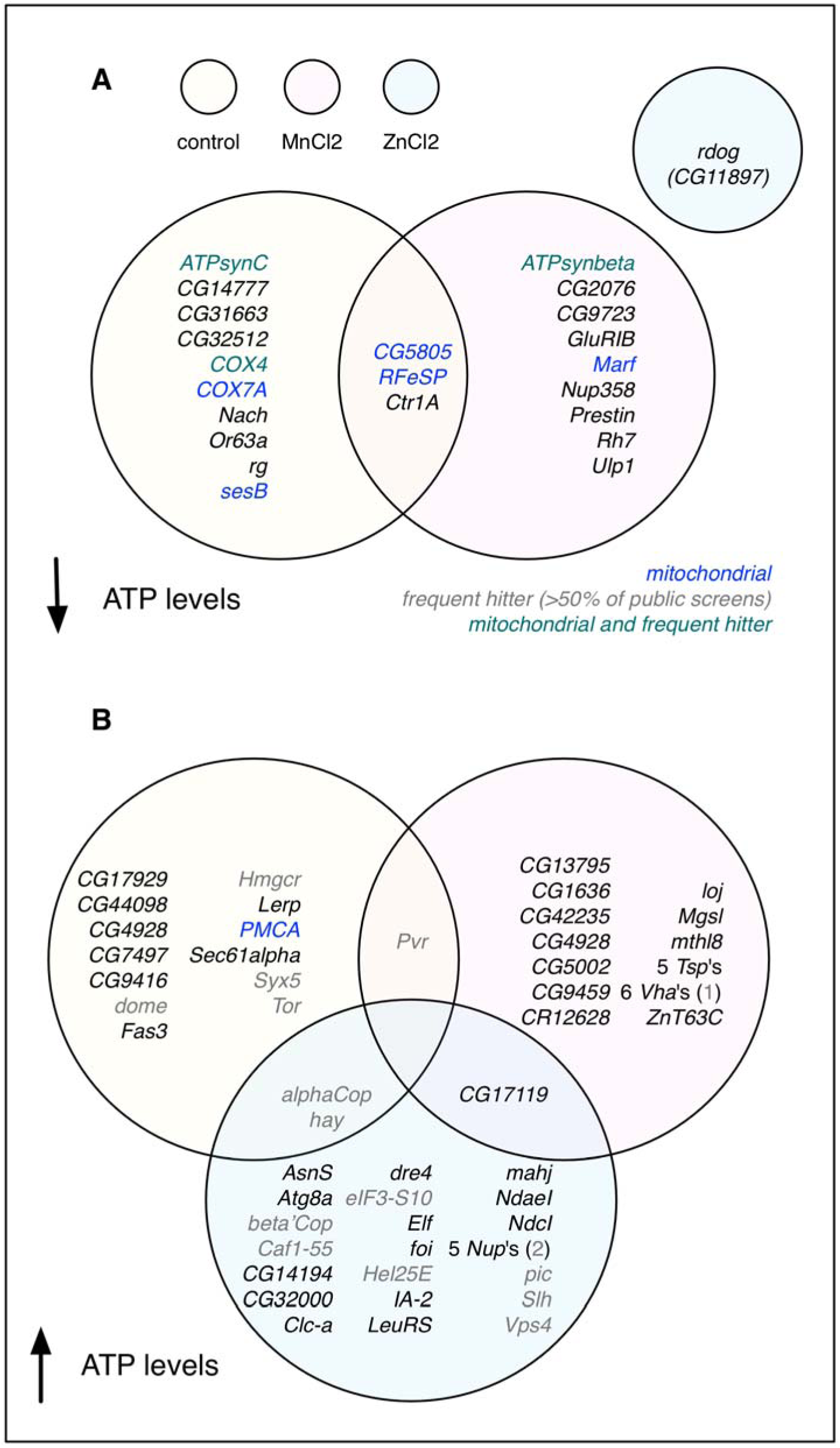
RNAi screen hits for control and metal toxicity conditions. Yellow circles, control conditions; pink circles, MnCl_2_ supplemented conditions; blue circles, ZnCl_2_ supplemented conditions. Genes that encode mitochondrial components are shown in blue; genes that are ‘frequent hitters’ as defined by a percent hit rate of ≥ 50% of *Drosophila* screens in GenomeRNAi are shown in gray; genes that are both mitochondrial and frequent hitters are shown in forest green.

**Figure 2:**
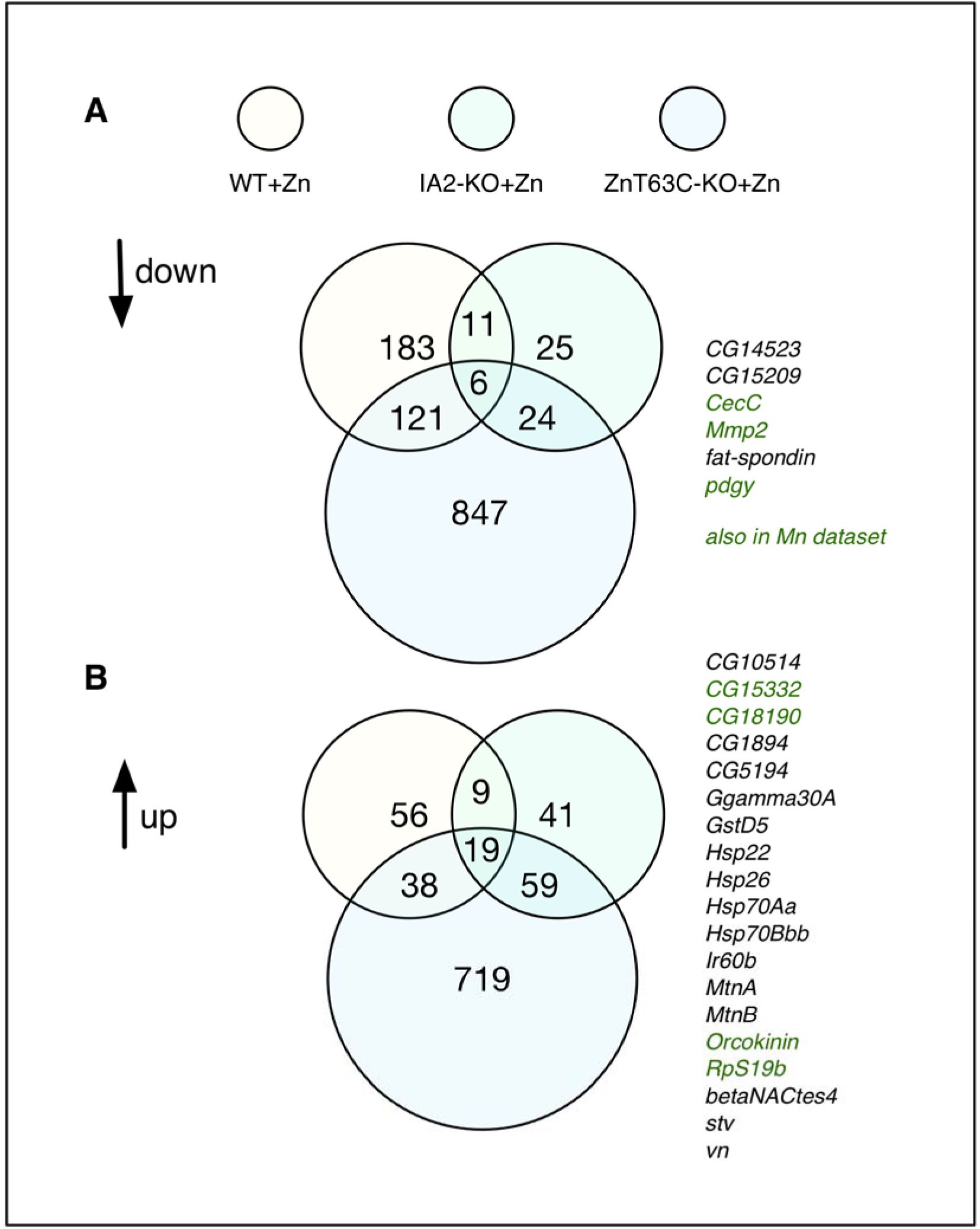
Overlap among genes down-or up-regulated in wildtype, *IA2-KO*, or *ZnT63C-KO* cells supplemented with zinc chloride. Yellow circles, wildtype (WT) zinc-treated cells; mint green circles, *IA2-KO* zinc-treated cells; blue circles, *ZnT63C-KO* zinc-treated cells. The six down-regulated and 19 up-regulated genes in common are shown to the right. Green text indicates genes also found in one or more genotype under manganese-treated conditions.

**Figure 3:**
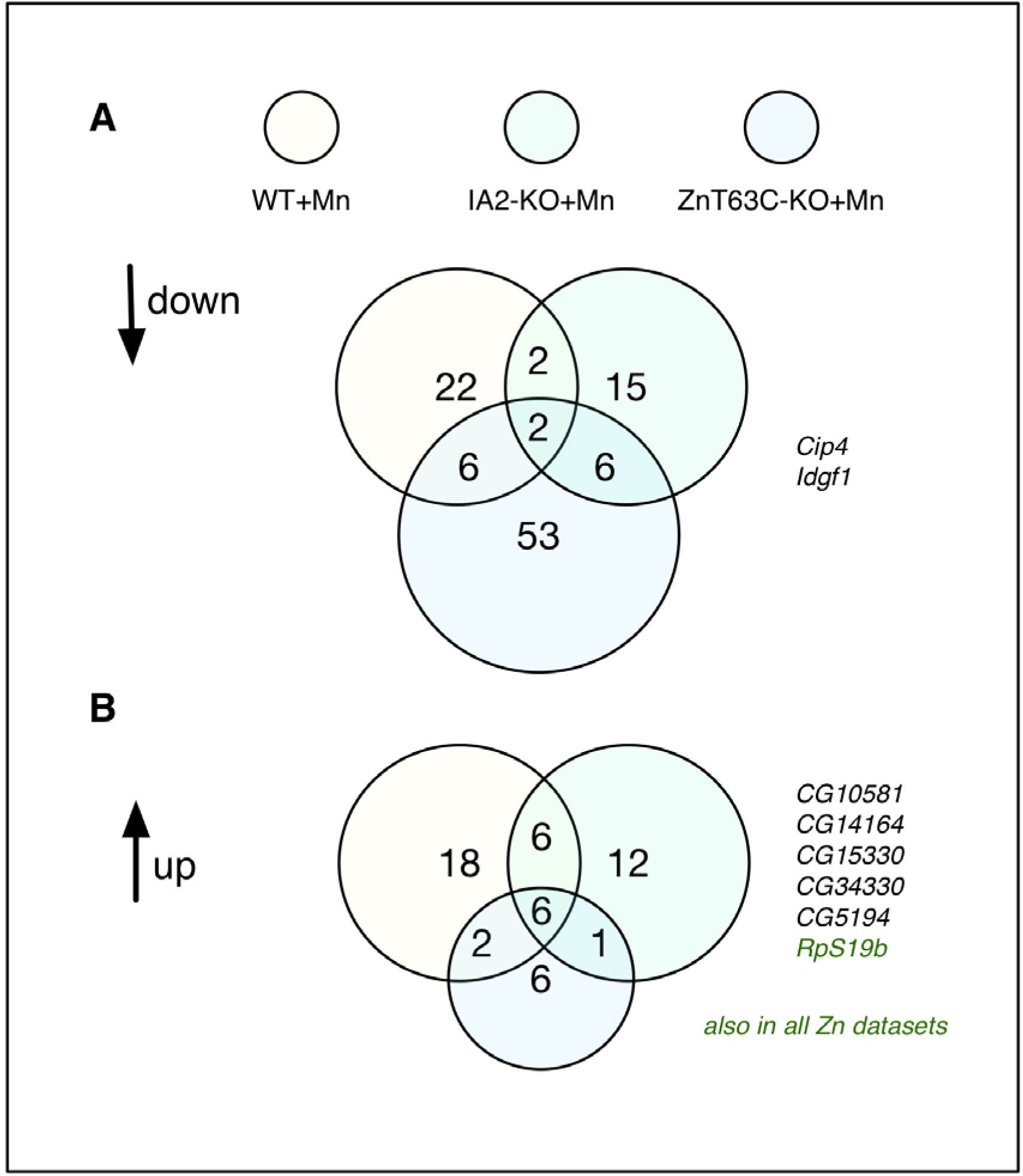
Overlap among genes down- or up-regulated in wildtype, *IA2-KO,* or *ZnT63C-KO* cells supplemented with manganese chloride. Yellow circles, wildtype (WT) manganese-treated cells; mint green circles, *IA2-KO* manganese -treated cells; blue circles, *ZnT63C-KO* manganese -treated cells. The two down-regulated and six up-regulated genes in common are shown to the right. Forest green text indicates genes a gene found in all three genotypes under zinc-treated conditions.

## RESULTS AND DISCUSSION

### *Drosophila* cell-based screen for modifiers of metal chloride toxicity

We reasoned that the application of high-throughput ‘omics approaches in *Drosophila* cultured cells would provide a robust dataset that can inform future *in vivo* studies. To get a genome-scale view of genes relevant to zinc detoxification, we performed three large-scale RNA interference (RNAi) screens in parallel using *Drosophila* S2R+ cultured cells. We used total ATP levels as an indirect assay readout of cell viability and number, and screened under three conditions: control conditions, toxic zinc conditions (i.e. supplementation to a concentration of 15 mM ZnCl_2_), and toxic manganese conditions (15 mM MnCl_2_). Although our main goal was to identify genes relevant to zinc, we chose to screen in parallel under toxic manganese conditions for two reasons. First, we reasoned that screening with an equimolar concentration of a different metal chloride salt would help exclude the possibility that the genes we identify in the zinc intoxication screen are relevant to the chloride ion, changes in osmolarity, or other non-specific effects. Second, we reasoned that the data would provide a helpful high-throughput dataset relevant to the biology of manganese, a metal that, like zinc, is essential for cell viability and toxic in excess but underexplored in the literature.

In order to quickly identify high-confidence screen ‘hits’ (positive results) at the gene level, we screened RNAi reagent libraries from the *Drosophila* RNAi Screening Center (DRSC) collection that have multiple unique RNAi reagents per gene coverage and optimized layouts. Specifically, we screened four libraries of double-stranded RNAs (dsRNAs): a large library targeting known or predicted *Drosophila* transmembrane domain-containing protein-encoding genes and three smaller libraries targeting genes encoding putative autophagy-related factors (ZnCl_2_ conditions only), conserved components of membrane-bound organelles, or candidate metal-related factors (total of 3885 dsRNAs targeting 1960 unique genes). Each screen condition was internally normalized (see Materials and Methods). How to access screen data is described in the data availability section of the Materials and Methods.

Several indicators suggest that the screens resulted in high-quality, metal-specific results. First, consistent with identification of condition-specific results, replicate assay plates within a treatment condition correlate well (Pearson correlation coefficients of 0.88 - 0.93) but replicate assay plates from different treatment groups do not (Pearson correlation coefficients of 0.24 - 0.50). Second, for many genes, two unique reagents targeting the same gene scored as hits under the same conditions and in the same direction. We define this set genes for which two unique reagents scored in the direction in a given condition as the set of ‘high-confidence hits’ (Fig. 1 and Supplemental Tables 1, 2, and 3). Third, the set of high-confidence hits in any given condition and direction include multiple members of a protein family (e.g. several tetraspanin family proteins are high-confidence hits in the same direction in the manganese screen), multiple members of a protein complex (e.g. several nuclear pore components are high-confidence hits in the same direction in the zinc screen), or multiple components of a given organelle (e.g. for both control and manganese conditions, mitochondrial proteins are among the high-confidence hits with lower ATP levels vs. the internal controls; Fig. 1, blue text).

Altogether, we identified 30 high-confidence hits in the control treatment group, 29 in the zinc toxicity treatment group, and 36 in the manganese toxicity treatment group (Fig. 1 and Supplemental tables 1, 2, and 3). There is limited overlap among genes that scored as hits from each of the three screens. Three genes (*CG5805, Ctr1A, RFeSP*) are in common between screen hits for control conditions, decreased ATP (“down”) direction and MnCl_2_ conditions, down direction (Fig. 1A). Two genes (*alphaCOP, hay*) are in common between screen hits for control conditions, increased ATP (“up”) direction and ZnCl_2_-supplemented conditions, up direction; one gene (*Pvr*) is in common between screen hits for control conditions, up direction and MnCl_2_-supplemented conditions, up direction; and one gene (*CG17119*) is in common between screen hits for ZnCl_2_-supplemented conditions, up direction and MnCl_2_-supplemented conditions, up direction (Fig. 1B). For the zinc toxicity screen, there was a clear bias in detection of high-confidence hits conferring higher ATP levels (28 high-confidence hits) as compared with lower ATP levels (1 high-confidence hit). We suspect that the zinc treatment conditions were so toxic it was difficult to detect a significant further reduction in total ATP levels vs. the internal control. In addition, some genes identified in the screens are ‘frequent hitters’ (Fig. 1, gray or green text), which we define here as hits scoring in >50% of *Drosophila* RNAi screens in the GenomeRNAi database (Schmidt *et al.* 2013).

### Analysis of zinc screen results

We found a number of zinc-related genes among the high-confidence hits in the zinc screen. For example, *fear of intimacy (foi)*, which encodes a ZIP family zinc influx protein (Mathews *et al.* 2005), was identified as a high-confidence hit for increased total ATP levels (“up” hits) in the zinc screen but not in the other screens (Table 3). The zinc screen hits in the up direction also include *CG32000*, a putative ortholog of human ATP13A2 (PARK9); evidence from mammalian cells suggests a role for ATP13A2 in lysosomal zinc transport (Kong *et al.* 2014; Park *et al.* 2014; Tsunemi and Krainc 2014). The zinc screen hits in this direction further include *IA-2 protein tyrosine phosphatase (IA-2),* an ortholog of human PTPRN (better known as IA2), which, like the human zinc transporter ZNT8, is a common autoantigen associated with type 1 diabetes (Arvan *et al.* 2012).

The single high-confidence hit in the lower ATP levels direction in the zinc screen (i.e. knockdown appears to confer zinc sensitivity; “down” hits) is a gene with the systematic name *CG11897* that we rename *red dog mine (rdog)* after the large Alaskan zinc mine of that name. The *rdog* gene encodes a member of the ABCC sub-family of ABC transporters that also includes yeast YCF1. Five additional genes were initially categorized as low-confidence hits in the ‘zinc treatment, lower ATP levels’ (down) category: *CG7627, CG3790, COX7AL, CR43469,* and *mthl3* (Supplemental file 1). Notably, like *rdog, CG7627* also encodes an ABCC family member, further supporting the idea that ABCC-type transporters are relevant to zinc chloride detoxification. Based on enrichment analysis (below), *CG7627* was promoted to a ‘moderate’ confidence hit (Supplemental file 1).

We performed enrichment analysis for gene ontology (GO) ‘cellular compartment’ or ‘biological process’ terms; pathways as annotated in Reactome (Croft *et al.* 2011); and protein complexes as annotated by COMPLEAT (Vinayagam *et al.* 2013). In all cases, we used the full set of hits in the analysis (i.e. both low- and high-confidence hits). We re-annotated low-confidence hits as ‘moderate’ confidence if they were members of a significant enrichment group (Supplemental file 1). Overall, the results of the enrichment analysis further suggest the quality and specificity of the screens. For the zinc toxicity screen, enrichment is driven by the presence of multiple components of the nuclear pore complex, suggesting the possible involvement of nuclear transport in zinc-induced cell death or another relevant process. We also note that there are related genes in the human genome for all of the high-confidence hits identified in the zinc screen (Supplemental Table 2).

### Transcriptomics analysis of wildtype and zinc-sensitized cells

The use of genetically zinc-sensitized strains of *Drosophila* has helped uncover mechanisms of zinc homeostasis *in vivo* (Lye *et al.* 2012; Lye *et al.* 2013). We reasoned that production of mutant cell lines lacking activity of the zinc exporter *ZnT63C* would similarly result in cultured cells genetically sensitized to zinc supplementation and allow for detection of zinc-related genes. Such an approach would allow us to capitalize on the advantages of performing transcriptomics studies in a relatively homogenous cultured cell line, allowing for robust detection of down- or up-regulated genes, as well as minimize the need to treat cells with high ionic strength solutions. We used a CRISPR-Cas9 strategy to target *ZnT63C,* which encodes a zinc efflux protein, and also targeted *IA-2*, which encodes the *Drosophila* ortholog of human PTPRN/IA2, which, like the human zinc transporter ZNT8, is a common autoantigen in type 1 diabetes (Arvan *et al.* 2012). Following transfection of CRISPR reagents, single-cell isolation, and identification of candidate knockout cell lines (see Materials and Methods), we confirmed by Sanger sequencing the presence of frameshift mutations in all copies of the genes in clonal cell lines. We will refer to the cell lines hereafter as *ZnT63-KO* and *IA2-KO*.

We next treated wildtype S2R+ cells, *IA2-KO*, or *ZnT63-KO* cells cells with mild zinc or manganese chloride supplementation, and isolated RNA from the samples for next-generation transcriptome sequencing (RNAseq). In total, we performed RNAseq analysis on two replicates each of nine combinations of genetic and treatment conditions: wildtype, *IA2-KO,* or *ZnT63C-KO* cells under control, mild zinc supplementation (1 mM ZnCl_2_), or mild manganese supplementation conditions (1 mM MnCl_2_). Following sequencing, we obtained analyzed ‘fragments per kilobase of transcript per million mapped reads’ (FPKM) values and used these as the dataset for all subsequent analyses. The number of down- or up-regulated genes for mild zinc and manganese chloride supplementation conditions, relative to values for untreated cells of the same genotype, is summarized in Table 1. We observed a larger overall transcriptional response in *ZnT63C-KO* cells treated with ZnCl_2_ than in the other genotypes and conditions (i.e. a larger total number of genes with significant log_2_ values as compared to the control and a large fold-change among the top hits), consistent with the idea that *ZnT63C-KO* are more sensitive to zinc supplementation than wildtype S2R+ cells. Supplemental file 2 includes a list of the genes the data suggest are down-or up-regulated in each genotype and condition. How to access the RNAseq data is outlined in the data availability section of the Materials and Methods.

As summarized in Table 1, we identified >1800 putative zinc-responsive genes. Several indicators suggest that the strategy of genetically sensitizing cells to zinc supplementation was successful in identifying high-confidence zinc-responsive genes, including the observation that several genes down- or up-regulated in two or more zinc-supplemented S2R+, *IA2-KO*, and/or *ZnT63C-KO* genotypes (Fig. 2). Not surprisingly, the set of zinc-responsive genes include metallothionines, which act as metal chelators, as well as a number of heat shock proteins (Fig. 2B). Interestingly, at least for some genes, the level of up-regulation induced by zinc in *IA2-KO* cells is intermediate as compared with wildtype and *ZnT63C-KO* zinc-treated cells (Supplemental Table 4, Supplemental file 2). As compared with zinc, supplementation with low levels of MnCl_2_ of any genotype resulted in a small number of genes showing significant down- or up-regulation as compared with wildtype untreated cells (see Table 1, Fig. 3 and Supplemental file 2), consistent with the idea that knockout of *ZnT63C* sensitizes cells to zinc but not manganese. Despite the smaller numbers, however, some genes are in common between the zinc and manganese datasets (Fig. 2 and 3, green text, and Supplemental file 2), suggesting that these genes might be chloride-responsive or general factors.

### Enrichment analysis of transcriptomics data from zinc-treated cells

As for the RNAi screen data, we performed enrichment analysis of the transcriptomics data to detect GO terms, molecular pathways, or protein complexes significantly enriched in these datasets (Supplemental file 2). Enrichment among genes down-regulated in response to ZnCl_2_ supplementation includes processes or protein complexes related to ribosomes (e.g. Reactome “translation,” p-value 6.88 x 10^-15^ in zinc-treated wildtype cells), and pathways or cellular components of respiratory electron transport. Enrichment among genes up-regulated in response to ZnCl_2_ supplementation includes processes or complexes involving heat-shock proteins and *starvin* (*stv),* an ortholog of the human “BCL2 associated athanogene 3” or BAG3 gene. Significant enrichment was also seen in zinc-treated wildtype cells for the GO molecular function heme-copper terminal oxidase activity; the 3 of 21 genes in this group that result in enrichment are *COX4L* and *COX7AL,* which encode subunits of cytochrome c oxidase, and *CG42376,* which encodes an ortholog of human “cytochrome c oxidase assembly factor 6” or COA6. For zinc-treated *ZnT63C-KO* cells, up-regulated genes are also enriched for glutathione-related activities (e.g. KEGG “glutathione metabolism,” p-value 3.62 x 10^-9^ in *ZnT63-KO* cells). Indeed, 9 of 11 GstD sub-family genes, other glutathione S-transferase genes, and *Glutamate-cysteine ligase catalytic subunit (Gclc)*, which is a rate-limiting enzyme in the glutathione synthesis pathway, are up-regulated in zinc-treated *ZnT63C-KO* cells.

Altogether, the transcriptomics data show that zinc stress results in up-regulation of metal chelators and heat-shock proteins, and suggests that zinc stress has specific impacts on mitochondrial function that elicit compensatory transcriptional responses. Moreover, the data obtained using a zinc-sensitized genotype suggest that under high zinc stress conditions, there is a significant need for conjugation of substrates to GSH. This is consistent with a recent report that glutathione S-transferase activity is relevant to methyl mercury toxicity in *Drosophila* (Vorojeikina *et al.* 2017) and with results obtained for other species that associate metal detoxification with GSH conjugation and/or flux (Perego and Howell 1997; Penninckx 2002; Gharieb and Gadd 2004; Nagy *et al.* 2006; Franchi *et al.* 2012; Seth *et al.* 2012).

### Comparison of functional screen and transcriptomics data

We reasoned that genes encoding proteins normally involved in zinc influx would be expected to score in the ‘up’ direction in the screen (higher ATP values as compared with the internal control, consistent with resistance to zinc treatment) and ‘down’ in response to zinc supplementation in the transcriptomics analysis. Consistent with this, we found that the high-confidence screen hit *foi* was down-regulated in zinc-treated S2R+ and zinc-treated *IA2-KO* as compared with untreated cells of the same genotype, and down-regulated in zinc-treated *ZnT63C-KO* cells as compared with wildtype untreated cells. In addition, *CG32000* was down-regulated in zinc-treated *ZnT63C-KO* cells as compared with wildtype untreated cells. We also compared the data for components of the nuclear pore. The low-confidence screen hit *Nup107* was down-regulated in zinc-treated wildtype and zinc-treated *ZnT63C* cells as compared with genotype controls, and in zinc-treated *IA2-KO* cells as compared with untreated cells of the same genotype. In addition, the high-confidence hit *Nup93-1* was down-regulated in zinc-treated *ZnT63C-KO* cells as compared with the wildtype untreated control, and the additional nuclear pore component-encoding genes *Nup43, Nup44A, Nup50, Nup54,* and *Nup160* were down-regulated in *ZnT63C-KO* cells as compared with untreated cells of the same genotype.

We next explored the converse prediction: that genes encoding proteins protective against zinc intoxication would be expected to score in the ‘down’ direction in the screen and to be up-regulated in response to zinc supplementation. Despite the relatively small number of ‘down’ direction hits in the zinc screen (Fig. 1 and Supplemental File 1), we did find overlap between the RNAi ‘down’ direction hits and zinc-responsive gene lists. The low-confidence hit *COX7AL* was significantly up-regulated in zinc-treated S2R+ and zinc-treated *ZnT63C-KO* cells as compared with untreated genotype controls, and in all three as compared with untreated wildtype cells. In addition, the one high-confidence ‘down’ direction RNAi screen hit, *rdog*, scored as significantly up-regulated in zinc-treated *ZnT63C-KO* cells; the log_2_ values for *rdog* were 1.21 for zinc-treated *IA2-KO* cells and 3.53 for zinc-treated *ZnT63C-KO* cells as compared with genotype controls.

### *rdog* is up-regulated in response to zinc in *Drosophila* S2R+ cells

We further confirmed that *rdog* is up-regulated in response to zinc supplementation using a graded series of ZnCl_2_ to supplement the culture media of wildtype S2R+ cells followed by quantitative real-time PCR (qPCR), as shown in Fig. 3. As expected, under ZnCl_2_-supplemented conditions, levels of the metallothionine-encoding gene *MtnA* levels are up-regulated and levels of the zinc importer-encoding gene *foi* are down-regulated. Under the same conditions, the levels of *rdog* are up-regulated (Fig. 3A). Based on identification of *rdog* in the ZnCl_2_ but not MnCl_2_-supplemented screen and transcriptomics datasets, we suspected that the effect is zinc-specific, rather than being attributable to the chloride ion. To further test this experimentally, we performed qPCR analysis on wildtype S2R+ cells supplemented with ZnSO_4_. The trends for control and *rdog* transcript levels are similar (Fig. 3B), demonstrating that *rdog* expression is up-regulated by zinc in *Drosophila* S2R+ cells.

### Analysis of parallel studies using manganese chloride

As mentioned, we performed the RNAi screens and transcriptomics studies with MnCl_2_ in parallel to help distinguish zinc-specific factors from general factors, and to provide an additional metal intoxication-related data resource. For the MnCl_2_ toxicity screen, enrichment among genes conferring higher ATP levels upon knockdown is driven by the presence of multiple components of the vacuolar H+ ATP transport machinery (Fig. 1B and Supplemental Table 2). This suggests the possible relevance of proton transport to manganese-induced cell death or another related process. Several tetraspanin family proteins are also hits in the manganese screen. This is intriguing, as three tetraspanin family proteins were detected as co-regulated by trans-eQTLs following feeding of flies with lead (Pb) (Ruden *et al.* 2009), suggesting the possibility of a general role for tetraspanin family proteins in detection or responses to metals or metal-induced stress. Enrichment analysis of genes conferring lower ATP levels in the MnCl_2_ screen points to the relevance of mitochondria. With regards to transcriptomics analysis, we found that the results for MnCl_2_-treated samples were more useful as a comparison set for ZnCl_2_ treatment than for detection of Mn-specific factors. Overall, the fold-change values were modest and a relatively small number of genes surpassed the cutoff values; for example, only 23 genes were down-regulated and 33 up-regulated in MnCl_2_-treated wildtype cells (Table 2). Nevertheless, overlap between these and the zinc treatment group was observed (Fig. 2 and 3, Supplemental file 2), suggesting that the manganese treatment group provides a useful filter for further refinement of a list of candidate zinc-related factors.

### Concluding remarks

The functional genomics and transcriptomics data sets described here provide a genome-scale resource for the study of zinc biology in *Drosophila*. Despite the fact that we performed the RNAi screens under high metal supplementation conditions, we were able to identify factors known to be relevant to zinc homeostasis at physiological levels (i.e. *foi* and *CG32000*). This is consistent with known overlap between metal homeostasis and detoxification genes, and suggests the validity of the approach. In addition, despite assay bias, we were able to detect one high-confidence gene, *rdog*, for which knockdown of results in lower ATP values as compared with the internal control. We further found that *rdog,* an ortholog of yeast YCF1, is up-regulated in genetically zinc-sensitized cells following mild zinc supplementation. Identification of *rdog* in the cell-based screen, as well as identification in the transcriptomics data of *rdog*, *Gclc,* and genes encoding glutathione S-transferase family proteins (Saisawang *et al.* 2012), supports the idea that GSH is relevant to zinc detoxification in *Drosophila*. Moreover, the observation that another gene encoding ABCC family member, *CG7627*, was also identified as a zinc sensitivity screen hit in this work, together with the fact that a third *Drosophila* ABCC family member, dMRP, has previously been implicated in methylmercury toxicity (Prince *et al.* 2014), suggest a general role for ABCC-type transporters in metal detoxification in *Drosophila*. Altogether, we expect that the ‘omics data presented here will guide further research into the mechanisms underlying metal homeostasis and detoxification in *Drosophila* and other systems. For example, the data provide a focused set of candidates for *in vivo* analyses of wildtype and genetically metal-sensitized flies under normal or metal-supplemented conditions, as well as for *in vivo* analyses in fly models of human diseases such as diabetes or neurodegeneration.

## ACKNOWLEDGMENTS

We thank Richard Burke (Monash University, Melbourne, Australia), Juan Antonio Navarro Langa (Universität Regensburg, Regensburg, Germany), Fanis Missirlis (Center for Research and Advanced Studies of the National Polytechnic Institute, Mexico City, Mexico), and Daniela Zarnescu (University of Arizona) for helpful conversations. In addition, we thank Chiao-Lin Chen, Kevin Kim, Afroditi Petsakou, Donghui Yang-Zhou, and other members of the DRSC/TRiP and Perrimon lab for helpful input on the project. We also thank the NYU RNAi Core Facility for collaboration on production of the TM RNAi library. The DRSC is supported by NIH NIGMS R01 GM067761 (N.P., PI). This work was also supported in part by NIH NIEHS R21ES025615 (N.P., PI). In addition, S.E.M. is supported in part by the Dana-Farber/Harvard Cancer Center, which is supported in part by NCI Cancer Center Support Grant # NIH 5 P30 CA06516. N.P. is an investigator of the Howard Hughes Medical Institute.

## Supplemental Materials

**Supplemental file 1, included as an Excel file**

**Supplemental file 2, included as an Excel file**

**Supplemental Reagents File, included as an Excel file in the prescribed format**

**Figure 4:**
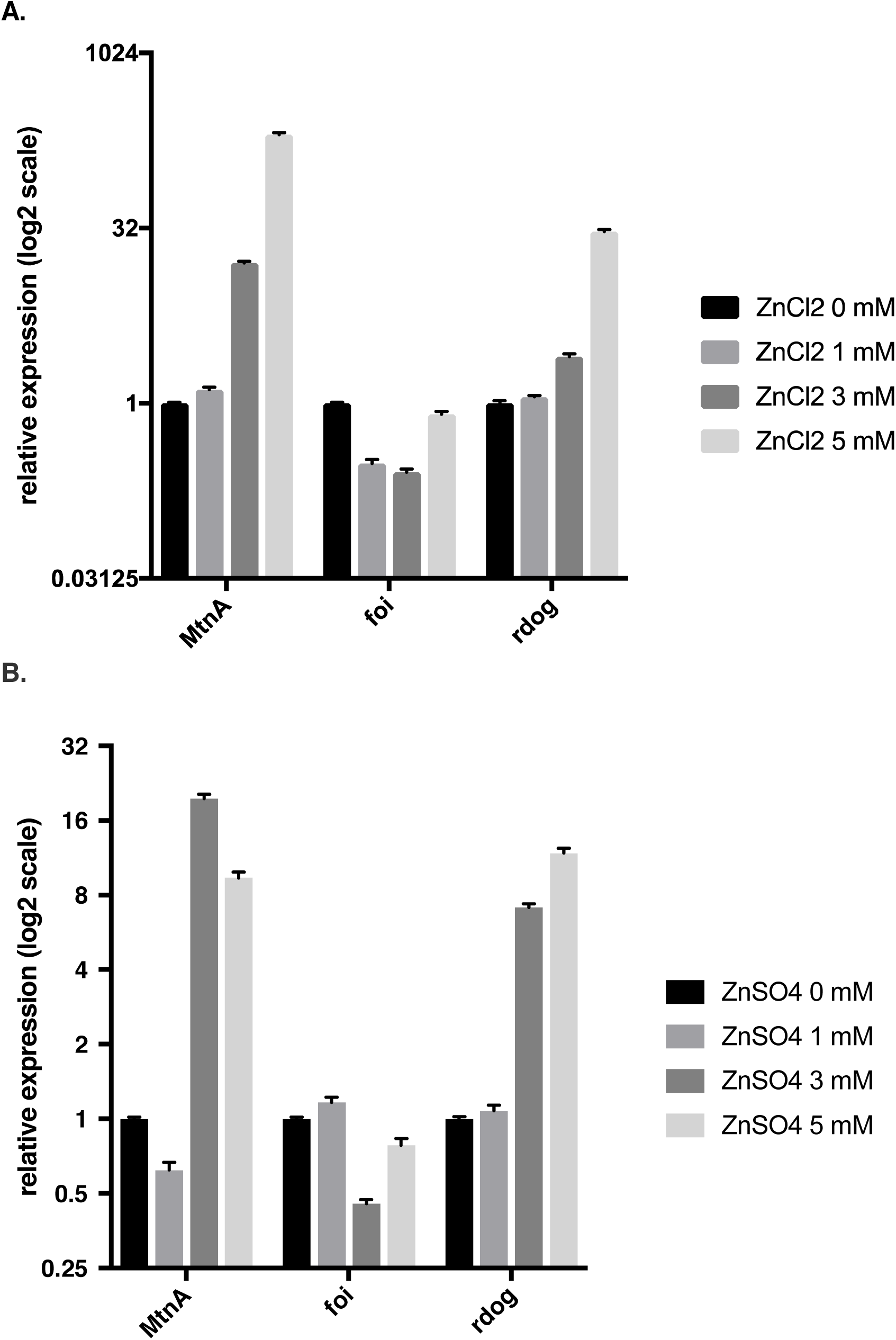
The ABCC-family transporter *rdog* is up-regulated in *Drosophila* S2R+ cells in response to zinc treatment. A, qPCR analysis of control and *rdog* transcript levels in S2R+ cells supplemented with ZnCl_2_ to a final concentration of 0, 1, 3, or 5 mM. The analyzed data are shown on a log_2_ scale. RNA levels from metal treatment samples were normalized to the 0 mM control (see Materials and Methods). Error bars indicate SEM. B, qPCR analysis of control and *rdog* transcript levels in S2R+ cells supplemented with ZnSO_4_ to a final concentration of 0, 1, 3, or 5 mM. The analyzed data are shown on a log_2_ scale. Consistent with a response to zinc, *rdog* transcript levels are up-regulated in response to zinc sulfate supplementation. RNA levels from metal treatment samples were normalized to the 0 mM control (see Materials and Methods). Error bars indicate SEM.

**Table S1:** High-confidence RNAi screen results for control cells

| FlyBase ID | Gene | Human | Treatment | Direction<sup>2</sup> | dsRNAs<sup>3</sup> | Avg Z |
| --- | --- | --- | --- | --- | --- | --- |
|  | symbol | ortholog<sup>1</sup> |  |  |  | score |
| FBgn0039830 | *ATPsynC* | ATP5G2 | Control | down | 2 | -2.17 |
| FBgn0026872 | *CG14777* | MPV17L | Control | down | 2 | -1.82 |
| FBgn0051663 | *CG31663* | -- | Control | down | 2 | -1.58 |
| FBgn0052512 | *CG32512* | TMEM205 | Control | down | 2 | -2.49 |
| FBgn0039223 | *CG5805* | SLC25A44 | Control | down | 2 | -1.97 |
| FBgn0032833 | *COX4* | COX4I1,2 | Control | down | 2 | -1.76 |
| FBgn0040529 | *COX7A* | COX7A2 | Control | down | 2 | -2.02 |
| FBgn0062413 | *Ctr1A* | SLC31A1 | Control | down | 2 | -2.06 |
| FBgn0262743 | *Fs(2)Ket* | KPNB1 | Control | down | 2 | -3.91 |
| FBgn0024319 | *Nach* | SCNN1B,G | Control | down | 2 | -1.89 |
| FBgn0035382 | *Or63a* | -- | Control | down | 2 | -1.73 |
| FBgn0021906 | *RFeSP* | UQCRFS1 | Control | down | 2 | -2.93 |
| FBgn0266098 | *rg* | NBEA | Control | down | 2 | -1.82 |
| FBgn0003360 | *sesB* | SLC25A4 | Control | down | 2 | -1.83 |
| FBgn0025725 | *alphaCOP* | COPA | Control | up | 2 | 2.11 |
| FBgn0038415 | CG17929 | -- | Control | up | 2 | 2.17 |
| FBgn0264907 | CG44098 | -- | Control | up | 2 | 1.76 |
| FBgn0027556 | CG4928 | UNC93A | Control | up | 2 | 1.99 |
| FBgn0036742 | CG7497 | PTGER1,3,4 | Control | up | 2 | 1.71 |
| FBgn0034438 | CG9416 | ERMP1 | Control | up | 2 | 1.85 |
| FBgn0043903 | dome | -- | Control | up | 2 | 1.82 |
| FBgn0000636 | Fas3 | -- | Control | up | 2 | 2.97 |
| FBgn0001179 | hay | ERCC3 | Control | up | 2 | 3.19 |
| FBgn0263782 | Hmgcr | HMGCR | Control | up | 2 | 2.59 |
| FBgn0051072 | Lerp | IGF2R | Control | up | 2 | 1.69 |
| FBgn0259214 | PMCA | ATP2B1,2,3,4 | Control | up | 2 | 2.29 |
| FBgn0032006 | Pvr | FLT1 | Control | up | 2 | 5.12 |
| FBgn0086357 | Sec61alpha | SEC61A1,2 | Control | up | 2 | 2.48 |
| FBgn0011708 | Syx5 | STX5 | Control | up | 3 | 2.87 |
| FBgn0021796 | Tor | MTOR | Control | up | 2 | 2.51 |

<sup>1</sup> Best match ortholog(s) are shown (DIOPT score cutoff >2) (Hu *et al.* 2011).
<sup>2</sup> Down, decreased total ATP levels following plate-based normalization within a treatment group; up, increased total ATP levels following plate-based normalization within a treatment group.
<sup>2</sup> Number of unique dsRNAs in the library that target the gene. For all high-confidence hits as shown here, each of the designs resulted in a Z-score >1.5 or <-1.5 and in the same direction as what was found for other designs targeting the same gene.

**Table S2:** High-confidence RNAi screen results for zinc chloride-treated cells

| **FlyBase ID** | **Gene**<br>**symbol** | **Human**<br>**ortholog(s)<sup>1</sup>** | **Treatment** | **Direction<sup>2</sup>** | **dsRNAs<sup>3</sup>** | **Avg Z**<br>**score** |
| --- | --- | --- | --- | --- | --- | --- |
| FBgn0039644 | *rdog* | ABCC family | ZnCl<sub>2</sub> | down | 2 | -1.81 |
| FBgn0025725 | *alphaCOP* | COPA | ZnCl<sub>2</sub> | up | 3 | 2.47 |
| FBgn0270926 | *AsnS* | ASNS | ZnCl<sub>2</sub> | up | 2 | 1.85 |
| FBgn0052672 | *Atg8a* | GABARAP | ZnCl<sub>2</sub> | up | 2 | 2.20 |
| FBgn0025724 | *beta'COP* | COPB2 | ZnCl<sub>2</sub> | up | 3 | 2.54 |
| FBgn0263979 | *Caf1-55* | RBBP4,7 | ZnCl<sub>2</sub> | up | 2 | 2.09 |
| FBgn0030996 | *CG14194* | TMEM185A,B | ZnCl<sub>2</sub> | up | 2 | 3.22 |
| FBgn0039045 | *CG17119* | CTNS | ZnCl<sub>2</sub> | up | 2 | 4.88 |
| FBgn0052000 | *CG32000* | ATP13A2,3,4,5 | ZnCl<sub>2</sub> | up | 3 | 4.14 |
| FBgn0051116 | *ClC-a* | CLCN1,2 | ZnCl<sub>2</sub> | up | 2 | 3.04 |
| FBgn0002183 | *dre4* | SUPT16H | ZnCl<sub>2</sub> | up | 3 | 3.30 |
| FBgn0037249 | *eIF3-S10* | EIF3A | ZnCl<sub>2</sub> | up | 3 | 2.22 |
| FBgn0020443 | *Elf* | GSPT1,2 | ZnCl<sub>2</sub> | up | 2 | 1.72 |
| FBgn0024236 | *foi* | SLC39A family | ZnCl<sub>2</sub> | up | 2 | 3.42 |
| FBgn0001179 | *hay* | ERCC3 | ZnCl<sub>2</sub> | up | 2 | 3.21 |
| FBgn0014189 | *Hel25E* | DDX39A,B | ZnCl<sub>2</sub> | up | 3 | 7.74 |
| FBgn0031294 | *IA-2* | PTPRN,N2 | ZnCl<sub>2</sub> | up | 2 | 2.54 |
| FBgn0284253 | *LeuRS* | LARS, LARS2 | ZnCl<sub>2</sub> | up | 2 | 1.73 |
| FBgn0034641 | *mahj* | DCAF1 | ZnCl<sub>2</sub> | up | 3 | 1.87 |
| FBgn0259111 | *Ndae1* | SLC4A family | ZnCl<sub>2</sub> | up | 2 | 1.60 |
| FBgn0039125 | *Ndc1* | NDC1 | ZnCl<sub>2</sub> | up | 2 | 1.87 |
| FBgn0039004 | *Nup133* | NUP133 | ZnCl<sub>2</sub> | up | 2 | 3.01 |
| FBgn0021761 | *Nup154* | NUP155 | ZnCl<sub>2</sub> | up | 3 | 2.69 |
| FBgn0039302 | *Nup358* | RANBP2 | ZnCl<sub>2</sub> | up | 2 | 3.17 |
| FBgn0027537 | *Nup93-1* | NUP93 | ZnCl<sub>2</sub> | up | 2 | 5.60 |
| FBgn0039120 | *Nup98-96* | NUP98 | ZnCl<sub>2</sub> | up | 2 | 6.02 |
| FBgn0260962 | *pic* | DDB1 | ZnCl<sub>2</sub> | up | 3 | 3.22 |
| FBgn0264978 | *Slh* | SCFD1 | ZnCl<sub>2</sub> | up | 2 | 1.58 |
| FBgn0283469 | *Vps4* | VPS4A,B | ZnCl<sub>2</sub> | up | 2 | 4.60 |

<sup>1</sup> Best match ortholog(s) or paralog families are shown (DIOPT score cutoff >2) (Hu *et al.* 2011).
<sup>2</sup> Down, decreased total ATP levels following plate-based normalization within a treatment group; up, increased total ATP levels following plate-based normalization within a treatment group.
<sup>2</sup> Number of unique dsRNAs in the library that target the gene. For all high-confidence hits as shown here, all of the designs resulted in a 'hit' as defined by a Z-score >1.5 or <-1.5 and in the same direction as what was found for other designs targeting the same gene.

**Table S3:**
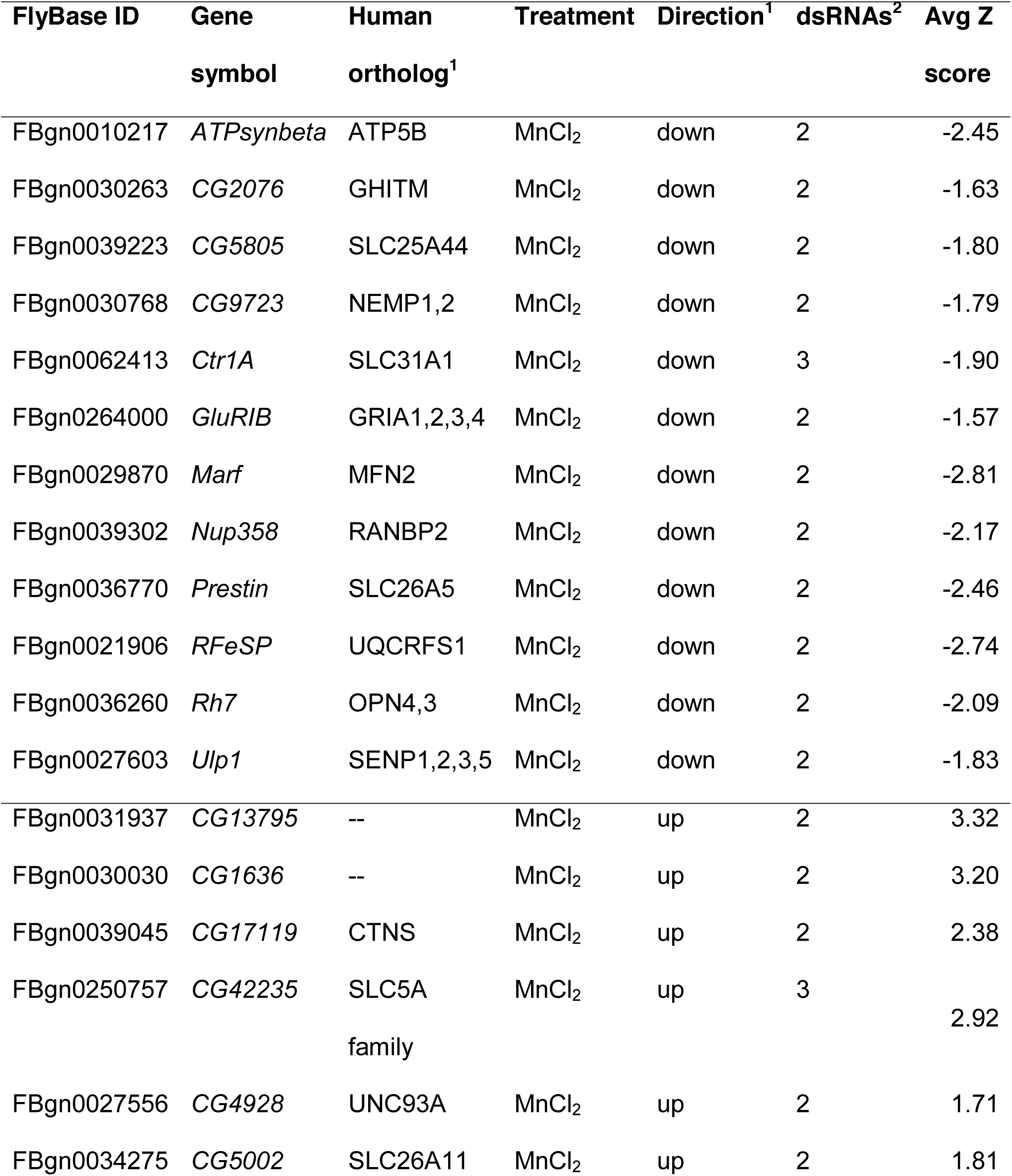
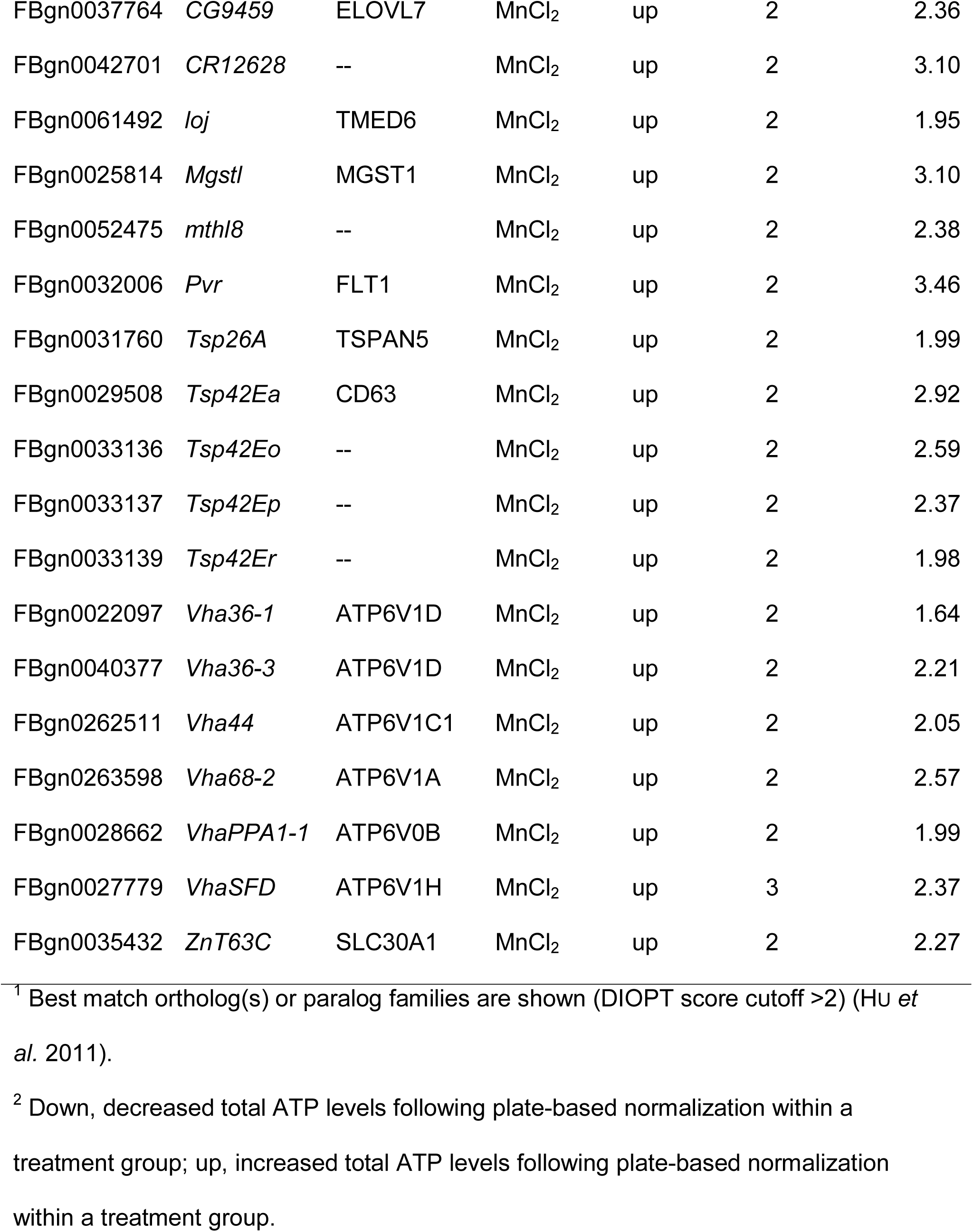
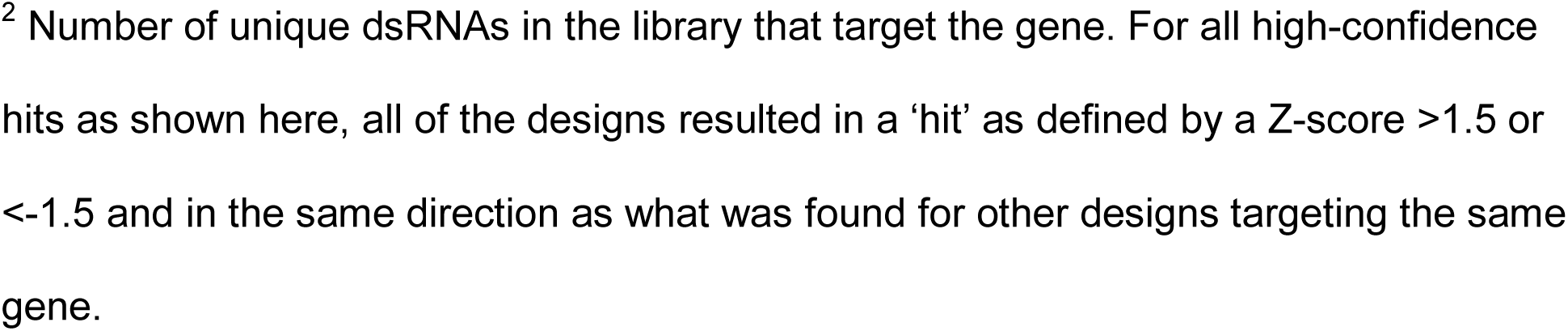
High-confidence RNAi screen results for manganese chloride-treated cells

**Table S4:** Genes highly up-regulated in response to zinc chloride supplementation of zinc-sensitized cells.

| **FlyBase ID** | **Gene** | **log2 WT+Zn/WT** | **log2 IA2+Zn/IA2** | **log2 ZnT+Zn/ZnT** |
| --- | --- | --- | --- | --- |
| FBgn0053192 | MtnD | -- | 3.63 | 9.47 |
| FBgn0002869 | MtnB | 1.73 | 4.21 | 8.93 |
| FBgn0267252 | Ggamma30A | 2.15 | 2.65 | 6.73 |
| FBgn0002868 | MtnA | 2.81 | 3.95 | 6.31 |
| FBgn0001223 | Hsp22 | 1.42 | 2.62 | 6.18 |
| FBgn0052274 | Drsl1 | -- | -- | 6.16 |
| FBgn0013277 | Hsp70Ba | -- | 3.57 | 6.05 |
| FBgn0051354 | Hsp70Bbb | -- | 3.62 | 5.99 |
| FBgn0013275 | Hsp70Aa | 1.86 | 2.41 | 5.97 |
| FBgn0010038 | GstD2 | -- | 3.40 | 5.92 |
| FBgn0013279 | Hsp70Bc | -- | 3.77 | 5.85 |
| FBgn0001225 | Hsp26 | 1.47 | 2.81 | 5.60 |
| FBgn0004635 | rho | -- | -- | 5.56 |
| FBgn0063493 | GstE7 | 1.43 | -- | 5.31 |
| FBgn0010041 | GstD5 | 1.56 | 2.06 | 5.24 |
| FBgn0031689 | Cyp28d1 | -- | -- | 5.24 |
| FBgn0015300 | Ssl | -- | -- | 5.15 |
| FBgn0013276 | Hsp70Ab | -- | 3.40 | 4.98 |
| FBgn0026320 | Tom | -- | -- | 4.94 |
| FBgn0004052 | Z600 | -- | -- | 4.90 |
| FBgn0001258 | ImpL3 | -- | 2.08 | 4.85 |
| FBgn0050413 | CG30413 | -- | 2.48 | 4.84 |
| FBgn0028940 | Cyp28a5 | -- | -- | 4.83 |
| FBgn0001230 | Hsp68 | -- | 2.87 | 4.78 |
| FBgn0259923 | Sep4 | -- | -- | 4.68 |
| FBgn0034741 | CG4269 | -- | -- | 4.60 |

All genes with a $\log_2$ >4.50 in zinc-treated *ZnT63C-KO* vs. untreated *ZnT63C-KO* cells are shown (see also Supplemental file 2). WT, wildtype; IA2, IA2-KO; ZnT, ZnT63C-KO; --, results not significant in this genotype.

